# Heterogenous transcriptomes of *Nicotiana tabacum* BY-2 suspension cell lines adapted to various osmoticum

**DOI:** 10.1101/2024.04.24.590873

**Authors:** Tomasz Skrzypczak, Przemysław Wojtaszek, Anna Kasprowicz-Maluśki

## Abstract

Plants abiotic stress response and adaptation belong to the most important subjects in plants biology. Here, we present *Nicotiana tabacum* suspension cell lines adapted during long term cultures to high concentrations of NaCl, KCl, mannitol and sorbitol. Obtained lines differ in osmotic stress agents and final media osmolarities. RNA-seq analysis revealed similarities, as well as differences in adapted lines transcriptomes. Presented here BY-2 cells lines form a good model to reveal molecular mechanisms of plants adaptations to salt and osmotic stress on cellular level.

## Introduction

Global crops cultures are severely affected by biotic and abiotic stresses, which often occur simultaneously or one after another. A combination of heat, drought and salt stress provide a danger, especially for irrigated farmlands in warm climate zones. Both, drought and salinity leads to osmotic stress. Long-term osmotic stress evokes activation of adaptive mechanisms, leading to changes in e.g. cellular metabolism, genes expression, mechanical adaptations^1–4^. Plants trigger response to perceived stress by transmitting inter-intracellular signaling, modifying phytohormones level and initiate transcriptional reprogramming. Four types of dehydration stress memory genes have been identified in *Arabidopsis thaliana*^5^. In response to stress plants inhibits growth and alter developmental processes, what contribute to growth-defense trade-off^6,7^. Additionally, plants could develop tolerance or adaptation to stress conditions, what require also transcriptional reprogramming ^4,8–10^. Molecular mechanisms of an abiotic stress memory that prime future stress and cross-stress responses were also revealed ^5,9,11,12^.

Impact of elevated [Na^+^][Cl^-^] was vastly investigated with focus on physiological, cellular and molecular aspects in *Arabidopsis thaliana,* crops, as well as in halophytes, which developed physiological and molecular adaptations to saline environment^2–4,13–26^. Despite growing risk of potassium overaccumulation in irrigated fields, there is not much information on plants cells response to high K^+^ concentration. However, it was shown that potassium toxicity surpasses sodium toxicity of the same concentration/water activity for *Arabidopsis thaliana*, due *to* its detrimental effect on nitrogen and phosphorus homeostasis ^27,28^. Excess of potassium and sodium, their specific and overlapping impact on physiology and metabolome was described also for *Vicia faba*^28^. In plants that synthesize mannitol and sorbitol, they play important roles in metabolism, osmoprotection and ROS quenching^29^. However, in high concentration in media they could leads to osmotic stress, induce water efflux and plasmolysis in cells from suspension culture.

*Nicotiana tabacum* BY-2 suspension cell lines are well established model in plants cell biology and biotechnology as well. BY-2 enables precise microscopy non-destructive observations correlated with reliable biochemical and molecular experiments^30^. It was shown that during adaptation of tobacco BY-2 suspension cells to the severe osmotic/salt stress significant modifications of extracellular polysaccharides (arabinogalactans and xyloglucans) and proteins content are occurring^31–33^. Adapted and non-adapted BY-2 cells differed also by cell walls rhamnogalacturonan, rhamnozyl, but not pectin level. Adapted cell lines were characterized also by thinner walls and smaller size^31,32^.

Research presented here shows, that after stepwise adaptation of tobacco BY-2 suspension cells to growth in the presence of concentrated osmotically active agents, BY-2 cell lines diverged into diverse, osmoticum type-specific lines. Such lines have been found as useful in research on specific molecular mechanisms of adaptations to long-term stress and cross-stress reactions. We characterized BY-2 cell lines morphology, and genes expression. Obtained phenotypic and transcriptome information about new tobacco suspension lines provide background for research on mechanisms of adaptation to stress, as well as cross-stress tolerance. Adapted BY-2 exhibit partially overlapping transcriptional changes, that elucidate osmoticum specific cells response.

## Abbreviations for BY-2 lines and respective RNA-seq samples

Control conditions BY-2 ->BY-2:Control

NaCl -> BY-2:NaCl

KCl -> BY-2:KCl

Mann -> BY-2:Mann

Sorb -> BY-2:Sorb

## Materials and methods

### Obtaining BY-2 cell lines

Tobacco BY-2 suspension cultured cells were maintained in liquid medium of pH 5.0, containing Murashige and Skoog salts^34^ and (per liter): 30 g sucrose, 100 mg inositol, 3 mg 2,4-D, 370 mg KH_2_PO_4_ and 1 mg thiamine-HCl, according to Nagata^35^. Cells were subcultured weekly at 10 ml of old culture per 70 ml of fresh medium in 300-mL Erlenmeyer flasks. Cultures were incubated at 21°C on a gyratory shaker at 120 rpm with a 2.5-cm displacement. During 18 months cells were gradually adapted to grow in medium supplemented with either 450 mM mannitol, 350 mM sorbitol, 140 mM KCl, or 190 mM NaCl. The cultures were transferred to fresh media containing increasing concentration of mannitol or NaCl in sequential manner, starting with 50 mM mannitol and 20 mM NaCl, but were otherwise handled identically to the control BY-2 cells. All of the results described herein are from cultures maintained in the presence of final concentrations of osmotic agents for more than 10 years, and all the sampling was done on the third day after transfer to fresh medium.

### BY-2 cells size and shape measurements

BY-2 cell lines were stained with 0.5% calcofluor white. No *a priori* cells features were utilized for measured morphology of living BY-2 cells. Cells (n = 50) images where taken with Nikon A1R confocal microscope, Plan Apo VC 60xA/1.2 WI; excitation 405 nm diode laser; detection 410-450 nm with PMT. Cells area were marked with polygon ROI. Fiji ImageJ was used to measure parameters: Area and Shape Descriptors.

### RNA isolation and RNA-seq analysis

RNA was quantified using a Qubit RNA Assay Kit (Life Technologies), and integrity was confirmed on an Agilent Bioanalyzer 2100 system. Library for Ilumina pair-ends sequencing was prepared with TruSeq Stranded mRNA LT Sample Prep Kit (Ilumina). Raw RNA-seq results from all 15 experiments were subjected to quality filtration by FASTQC^36^. HISAT2^37^was used for mapping to *N. tabacum* transcriptome (BioProject:319578; Ntab-TN90; Annotations: NCBI_100). StringTie^38^ was used for transcripts assembly, an expression profile was calculated with read counts and FPKM. Expression distribution after Relative Log Expression (RLE) was plotted and compared to raw data and log2 read counts. Differentially expressed genes (DEGs) were identified with nbinomWaldTest using DESeq2^39^ and for DEGs significance threshold conditions were |FC|>=2 and adjusted p-value<0.05. Expression heatmaps were made with bioinfokit^40^. Identified genes were functionally assigned to KEGG Pathways and BRITE hierarchy databases for *N. tabacum* genome (KEGG ID: T04643), as well as Gene Ontology with DAVID^41,42^ tool.

## Results

### Morphology of BY-2 suspension cell lines

Cultured BY-2 lines grow in concentrations of osmotica that were the highest in which culture were still growing after incremental increasing of concentrations. Increased osmolarity provide stress to cells structures, could lead to plasmolysis, and even death as well. Sodium and potassium ions induce also additional toxic effects, that fortify detrimental effects of increased osmolarity. Fresh media of analyzed lines differ by kind of osmotica, its final concentration, and consequently with final osmolarity, that ranged from ∼220 mOsm/kg for BY-2:Control to ∼750 mOsm/kg for BY-2:Mann. The intermediate osmolarities were measured for media for BY-2:Sorb (∼650 mOsm/kg), BY-2:KCl (∼500 mOsm/kg), and BY-2:NaCl (∼550 mOsm/kg).

BY-2:NaCl/KCl, and BY-2:Mann/Sorb lines show different morphologies and rate of cells aggregation (Figure 1). Elongated shape BY-2 cells in standard conditions grow in long string in which cells linked by shorter edges. In response to ionic agents (NaCl, KCl) cells increased their adhesive properties and formed clumps composed of randomly dividing cells. On the other hand, cells adapted to nonionic osmotica (mannitol, sorbitol) reveal more ordered pattern of precisely positioned cell divisions, resulting in the formation of files up to several tens of cells long. Lines that live in high salts consisted of smaller and less elongated cells. High salt BY-2 and BY-2:Sorb more rounded than BY-2:Control and BY-2:Mann. Moreover, BY-2:Mann/Sorb cells were averagely bigger than BY-2:Control. An increased osmolarity is known that leads to “softer” cell walls and cells swelling^43^. Additional ionic toxicity of NaCl/KCl might contribute to inhibition of cells growth and changes in division plane or cell walls adhesiveness, that result in cells clumping/aggregation.

**Figure 1.**
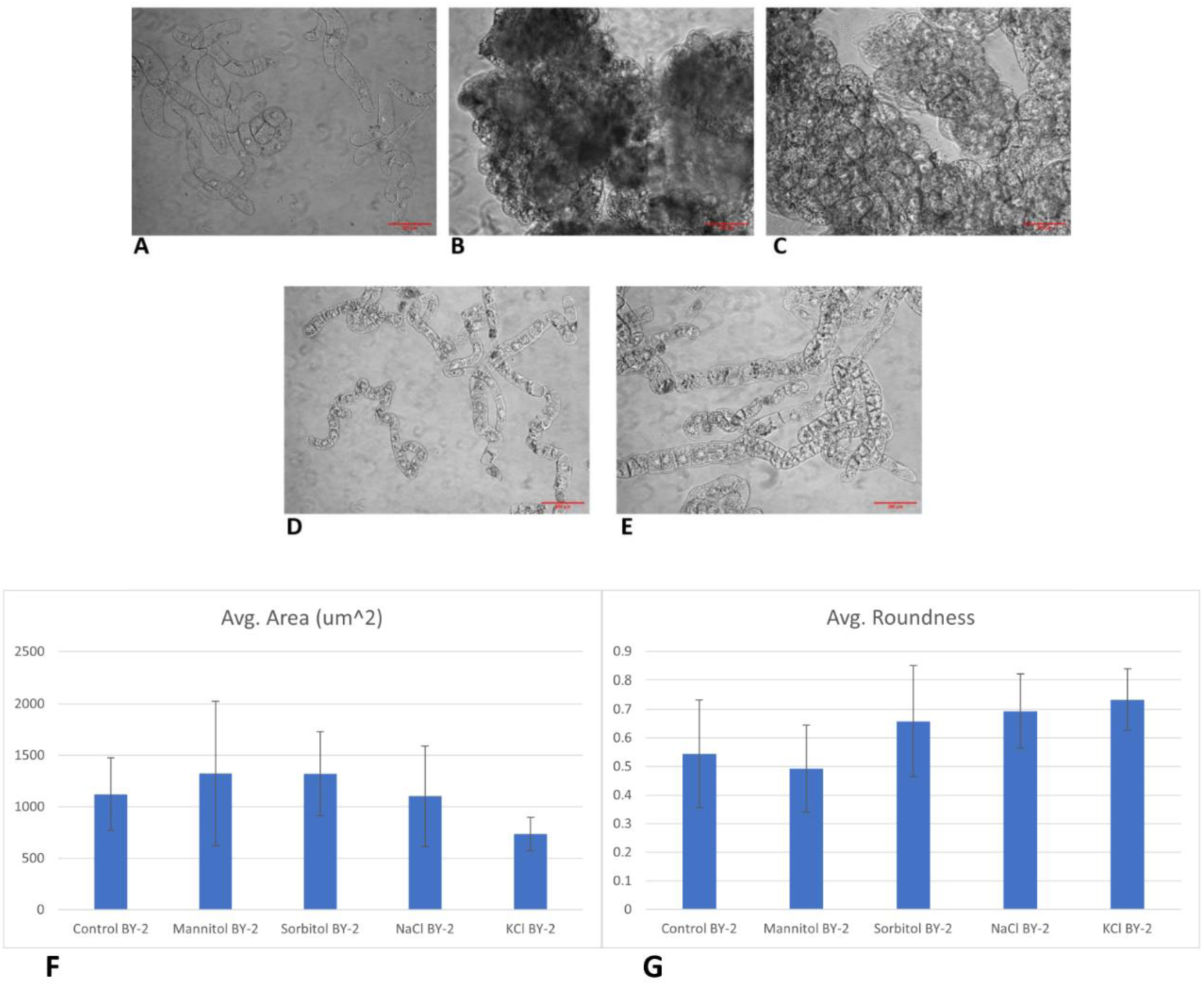
BY-2 suspension cell lines morphology: A) BY-2:Control; B) BY-2:NaCl; C) BY-2:KCl; D) BY-2:Mannitol; E) BY-2:Sorbitol. Live cells from lines with high salt concentrations strongly aggregate in cultures. F) Bar plot with average BY-2 cells area in um; G) Bar plot with average BY-2 cells roundness

### Differential Genes Expression analysis reveals differences between transcriptomes cell lines cultured in various osmoticum

In every sample >97% reads were mapped with Phred quality >20. All samples from certain lines occurred to be transcriptionally similar and consistent within replicates (data not shown). A consistency of expression within replicates was confirmed by hierarchical clustering and Multiple Component Analysis (MCA) of transcripts counts were used to identify similarity between transcriptomes and prove different gene expression patterns for every line (data not shown). By using each sample’s rlog transformed value, the expression similarities were clustered together in every BY-2 line for significant DEGs lists. Every BY-2 line was found to exhibit a clearly distinct gene expression pattern. Transcriptomes of BY-2:Mann and BY-2:Sorb that were the most similar to each other.

Similar amounts of DEGs around 2 000 were identified for every of adapted line paired with BY-2:Control. Numbers of DEGs ranged from 1874 for BY-2:KCl to 2338 for BY-2:Sorb (**Error! Reference source not found.**). DGE analysis confirmed the variabilities between genes up- and downregulated in BY-2:NaCl/KCl/Mann/Sorb (Figure 2). As many as approx. ¼ of DEGs, i.e. 444 genes, were common for all adapted lines paired with BY-2:Control. The common DEGs belong the most numerously to such KEGG pathways, like: Phenylpropanoid biosynthesis, MAPK signaling pathway - plant, Plant hormone signal transduction. The highest number of common DEGs for paired lines was found for BY-2:Mann and BY-2:Sorb (Fig. 6D). DEGs unique for only one line were the most numerous for BY-2:NaCl (886) and the least for BY-2:Mann (152). Total number of upregulated DEGs dominated over downregulated DEGs in all lines, except BY-2:KCl (Figure 2).

**Figure 2.**
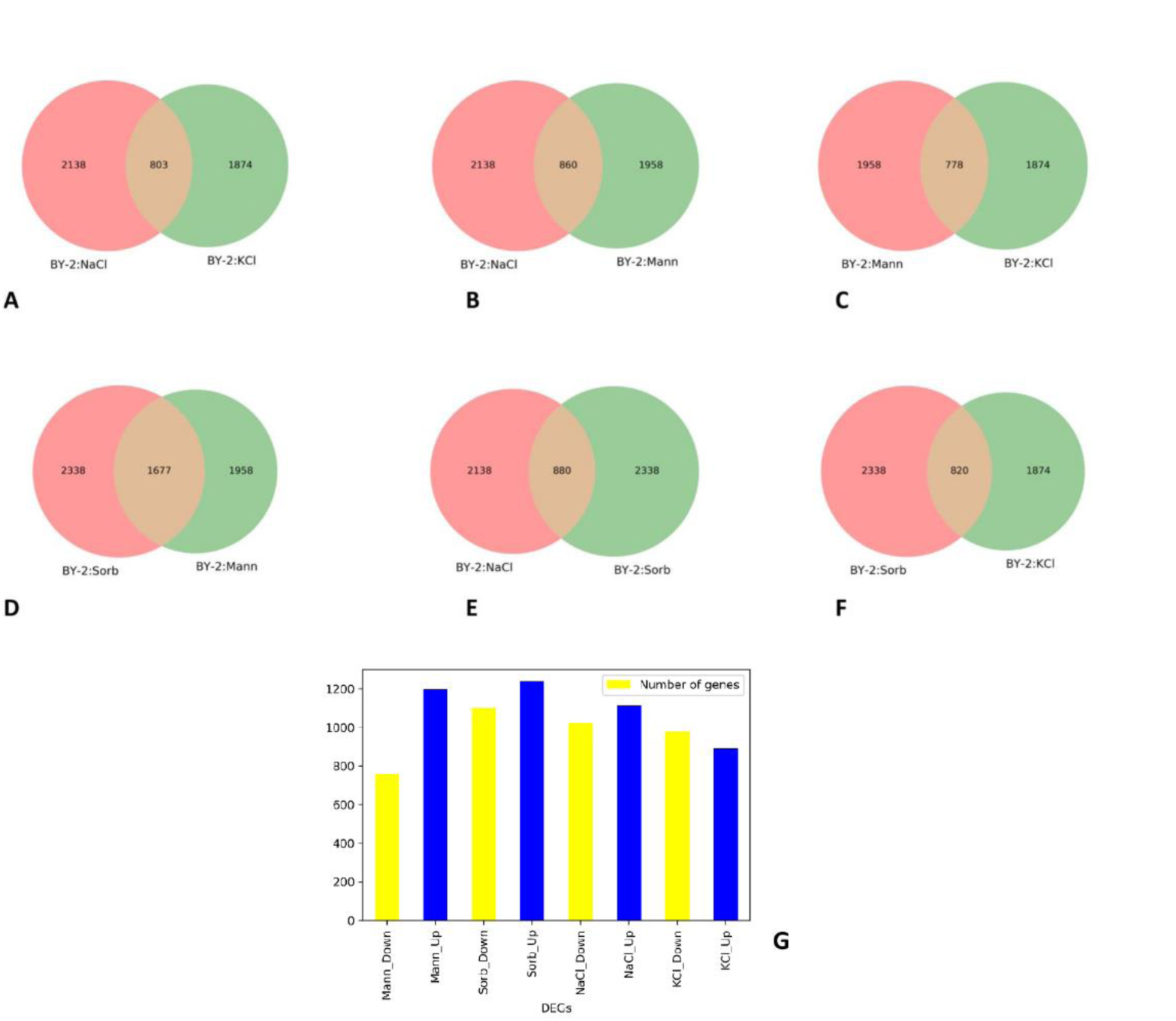
A) – F) Venn diagrams of DEGs numbers for BY-2 lines. G) number of significantly up- and downregulated genes in every BY-2 cell line. Uniquely, in the case of BY-2:KCl number of downregulated genes exceed upregulated genes.

### Gene expression functional analysis – *GO annotations*

DEGs for salts (BY-2:NaCl+KCl vs BY-2:Control), polyols (BY-2:Mann+Sorb vs BY-2:Control) adapted lines were assigned with Gene Ontology’s annotations. In both cases there were identified overrepresented genes with products annotated as integral membranes components (Figure 3 AB). Plasma membrane proteins, like channels and transporters play key role in maintaining osmotic and ionic homeostasis. Despite the fact that BY-2 cells as derived from root and do not contain developed chloroplasts, expression data suggest that plastids might maintain to be important organelles in adaptation of BY-2:Mann/Sorb to high osmotic pressure. Recently, there were reported plastids specialized in triggering stress signaling that shape^44,45^.

**Figure 3.**
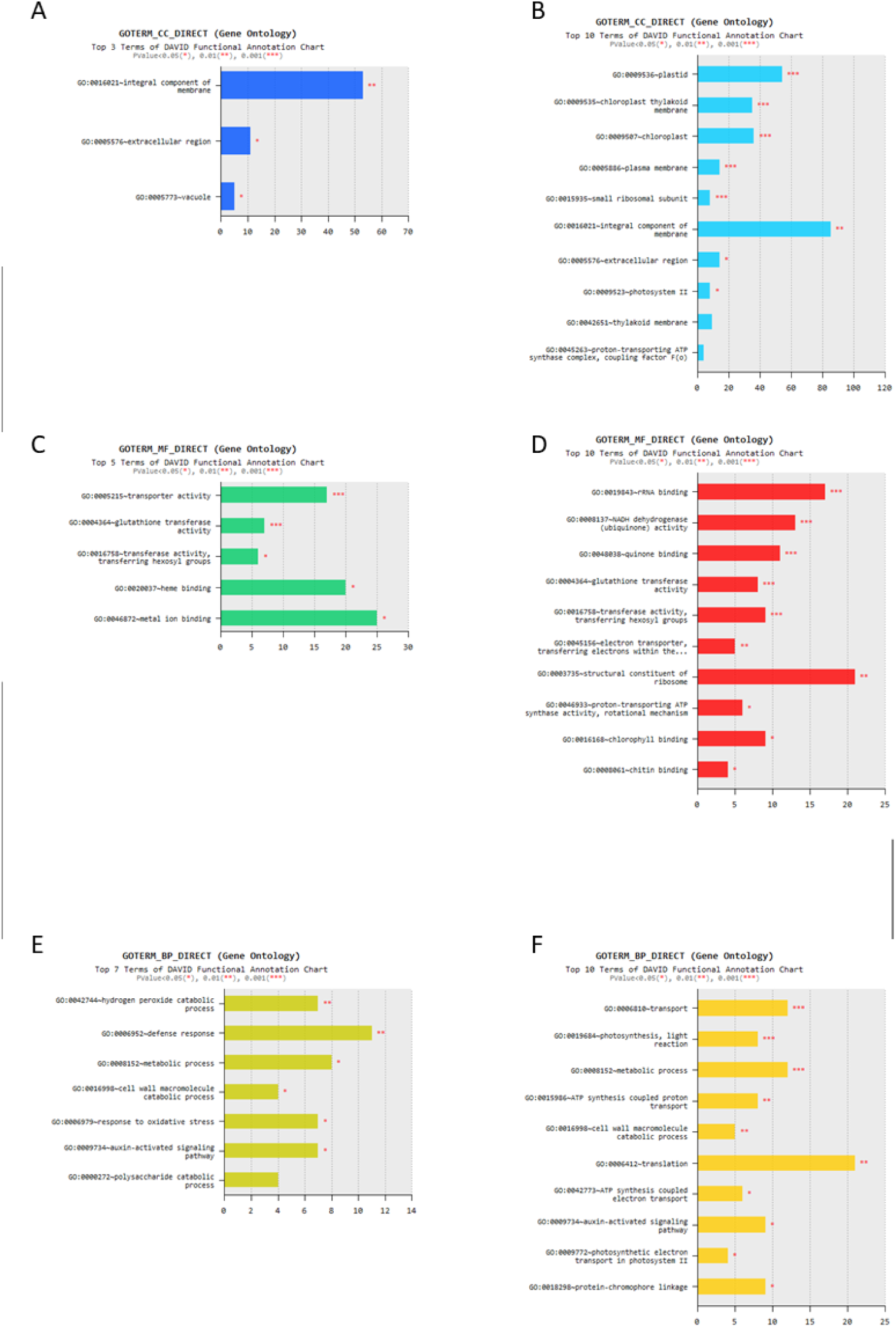
Gene Ontology categories overrepresented in DEGs of high salts (A, C, E) and high polyols (B, D, F) BY-2 lines

According to GO BP, DEGs from BY-2:NaCl/KCl, but not BY-2:Mann/Sorb contain overrepresented genes involved in response to oxidative stress, and defense response (Figure 3 EF) as well. Interestingly, in the case of BY-2:Mann/Sorb prominent fraction of DEGs was related to translation (GO BP), ribosomes and rRNA binding (GO CC, GO MF) (Figure 3 DF). It might suggest more substantial impact of high osmotic pressure, than ionic stress with lower osmotic pressure, on mechanisms of proteins biosynthesis could be expected in BY-2. However, also polyols specific indirect metabolic interaction with translation could not be excluded.

### Analysis of DEGs assigned to KEGG pathways -- affected by salt and osmotic stress conditions

Assignment of identified DEGs to KEGG pathways revealed processes important for BY-2 stress adaptation. Numerosity and variability of altered pathways pointed out massive changes in cells physiology that facilitate them to survive in high osmotic pressure and ionic stress conditions. Enrichment of KEGG pathways suggest that BY-2 lines have to adapt their basic anabolic and catabolic processes, as well as phytohormones production and signaling pathways. Widespread and strong overrepresentation of genes belonging to phenylopropanoid biosynthesis pathway (nta:00940), glycerophopsholipids (nta:00564), and phosphatidylinositol signaling system (nta:04070) might suggest occurrence also of transcriptionally regulated remodeling of cell walls and plasma membrane compositions.

Both salt and drought stress induce abscisic acid (ABA) synthesis and activation. Alterations in gene expression related to biosynthesis and signaling pathways of ABA/ethylene are well known response to abiotic stress. Here, the BY-2 suspension cell lines were exposed to high salt/polyol for long time - multiple generations, what might trigger specific adaptation/tolerance molecular mechanisms.

Expression of PP2C coding genes was significantly boosted in all stress adapted BY-2 lines, compared to BY-2:Control. Interestingly, line specificities of upregulated PP2Cs and SnRKs were observed (Fig. 10 A). We identified PP2Cs and SnRKs, that were boosted only in polyol or salt lines or uniquely in KCl BY-2 line (Figure 4 AB). The expression both of ABA (PYR/PYL) and ethylene (ETR/ETS) receptors was downregulated in all lines, what might be an adaptation to hypothetically elevated ABA and ethylene level (Figure 4 CD). Phytohormones receptors repression in expression was more pronounced in BY-2:Mann/Sorb, than in BY-2:NaCl/KCl. PYR/PYLs downregulation and PP2Cs upregulation were regularly shown in early response to drought, salt and ABA treatment (reviewed ^8^). Interestingly, ABA and ethylene related genes (KEGG nta:04016) exhibit stronger refinement in BY-2:Mann/Sorb, than in BY-2:NaCl/KCl. Downstream ethylene signaling effector gene Chitinase B (ChiB) is strongly upregulated in all BY-2 lines.

**Figure 4.**
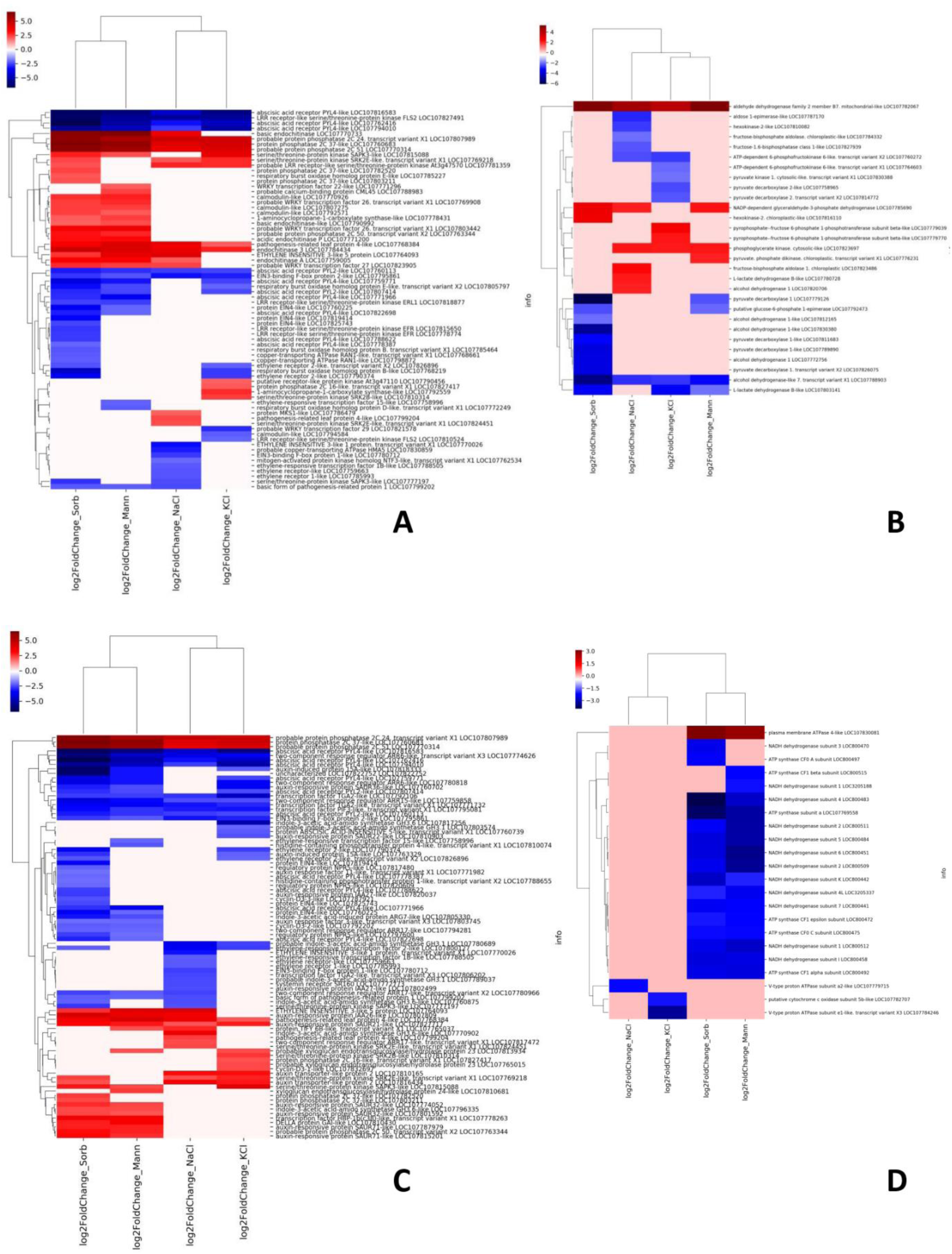
Heatmaps of log2 fold changes in gene expression. Genes derived from Nicotiana tabacum KEGG pathways: A) 04016 MAPK signalling pathway - plant; B) 00010 Glycolysis / Gluconeogenesis; C) 04075 Plant hormone signal transduction; D) 00190 Oxidative phosphorylation.

Genes expression data suggest that, ABA and ethylene levels in adapted BY-2 might be elevated, despite sustained high osmoticum concentrations for long time. High Na^+^, K^+^, Cl^-^ and high medium osmotic pressure induce ABA related stress response, even after multiple cells generations since concentrations had reached their final levels.

The reactive oxygen species (ROS) homeostasis has to be adjusted to specific environmental situation, what would enable cells to maintain ROS signaling and avoid ROS induced damages. In adapted BY-2, ROS producers coding genes – RboHs, and ROS scavengers coding genes (like CAT, APX, GST, CYP450) show line specific gene expression patterns (Figure 4 C,D,E; 5 A. 6 D). However, RboHs genes were extensively and commonly downregulated, what might prevent cells from producing too many ROS (Fig. 10A). In *Glycine max* it was shown that expression of ROS scavenging and producing genes were organ specific in the case of drought and heat combined response ^46^.

Interestingly, also genes involved in basic energy metabolism pathways, like glycolysis and oxidative phosphorylation, belong to the strongly changed pathways (Fig. 10BD). Prominent and the only one common for all four BY-2 lines was upregulation of aldehyde dehydrogenase (LOC107782067; mitochondrial-like ALDH2), that might be involved not only in glycolysis, but also in peroxidation of the poly-unsaturated fatty acids, or non-β-oxidative pathway of benzoic acid biosynthesis^47^. Alcohol dehydrogenase (LOC107788903) was commonly downregulated, what suggest lower oxygen consumption, and decreased risk of local hypoxia^48^. Interestingly, oxidative phosphorylation related genes, mostly NADH dehydrogenases and ATP synthases, were massively downregulated in BY-2:Mann/Sorb, but much less in BY-2:NaCl/KCl.

Lipid composition of plasma membrane play crucial role in biophysical properties, that provide basis of cells robustness and tolerance to higher osmotic pressure. Here, in BY-2 lines we observed numerous alterations of genes involved in glycophospholipids metabolism (nta:000564) (Fig. 6B). In maize, changes in glycophospholipids synthesis pathway’s genes expression were correlated with altered lipids composition after saline treatment^49^, and also lipodome alterations are expected in adapted BY-2.

Persistence of stress conditions requires from cells to permanently switch their transcriptional regulation network. Both up- and downregulated transcription factors were identified in BY-2 cell lines (Fig. 11A). TFs expression pattern provides key factors in regulation of transcriptome and cell’s biology in changed environmental conditions. Such TFs were found as differentially expressed like (Fig. 6 A):

1. In all lines downregulated were few genes coding bZIP TGA2-like, that are involved in salicylic acid dependent Systemic Acquired Resistance, but also in positive regulation of ROS homeostasis in response to UV-B^50–52^
2. Upregulation of WRKY TFs, which contribute to i.a. biotic and abiotic stress response in both ABA dependent and independent way^53–55^. Here, BY-2:KCl distinguishes from other lines with none upregulated WRKYs, and with one downregulated.
3. Downregulated were ETHYLENE INSENSITIVE-3 LIKE 1 and ethylene responsive transcription factors 2- and 15-Like, that might negatively regulate ethylene response^56–58^ . Oppositely ETHYLENE INSENSITIVE-3 LIKE 5 was commonly upregulated, however, its functions are elusive, even in *A. thaliana*.
4. Downregulation of two TCP7-like in BY-2:Sorb. These are involved in mediating environmental signals into growth responses, also during salt stress^59,60^
5. Upstream genes related to auxin signaling (like auxin transporter protein 1 - AUX1, AUX/IAA) were upregulated in all BY-2 lines. Significantly downstream, auxin responsive proteins, like GH3, AUX/IAA, SAUR families proteins show more diversified expression patterns within adapted lines.

Adapted BY-2 cell lines clearly differentiate their transcriptomes, also by regulation by active and available TFs. Some of the identified significant TFs (WRKY, EIL1, TCP7-like, TGA2-like) were also previously shown as induced/repressed in short-term response to stress. Many auxin, ABA, and ethylene related genes were commonly for both shrunken BY-2:NaCl/KCl and not shrunken BY-2:Mann/Sorb were identified. Hence, no clarifying impact of TFs or hormones related genes have been observed. Interestingly, BY-2:KCl shows the most distinctive pattern of TFs expression within adapted BY-2 lines.

It was reported that K^+^ in the same concentration as Na^+^ is more toxic to plants^27^. BY-2:KCl potential nitrogen starvation might be a reason for disturbed growth and morphology. It is known that Na^+^ tends to replace K^+^ in enzymatic reactions, so BY-2:NaCl and BY-2:KCl partially exhibits potassium deficiency and excess stress respectively^61^. However, here in BY-2:KCl almost no changes were observed in basic nitrogen metabolism (nta:00910) related genes, in opposite to BY-2:NaCl/Mann/Sorb, where differentially expressed genes were more abundant within nitrogen metabolism related genes (Fig. 5C). Moreover, also autophagy pathway was not enriched in DEGs in BY-2:KCl/NaCl DEGs (nta:04136; data not shown), despite it was shown in the case of *A. thaliana* and excess of Na^+^, as well as K^+19,27^

**Figure 5.**
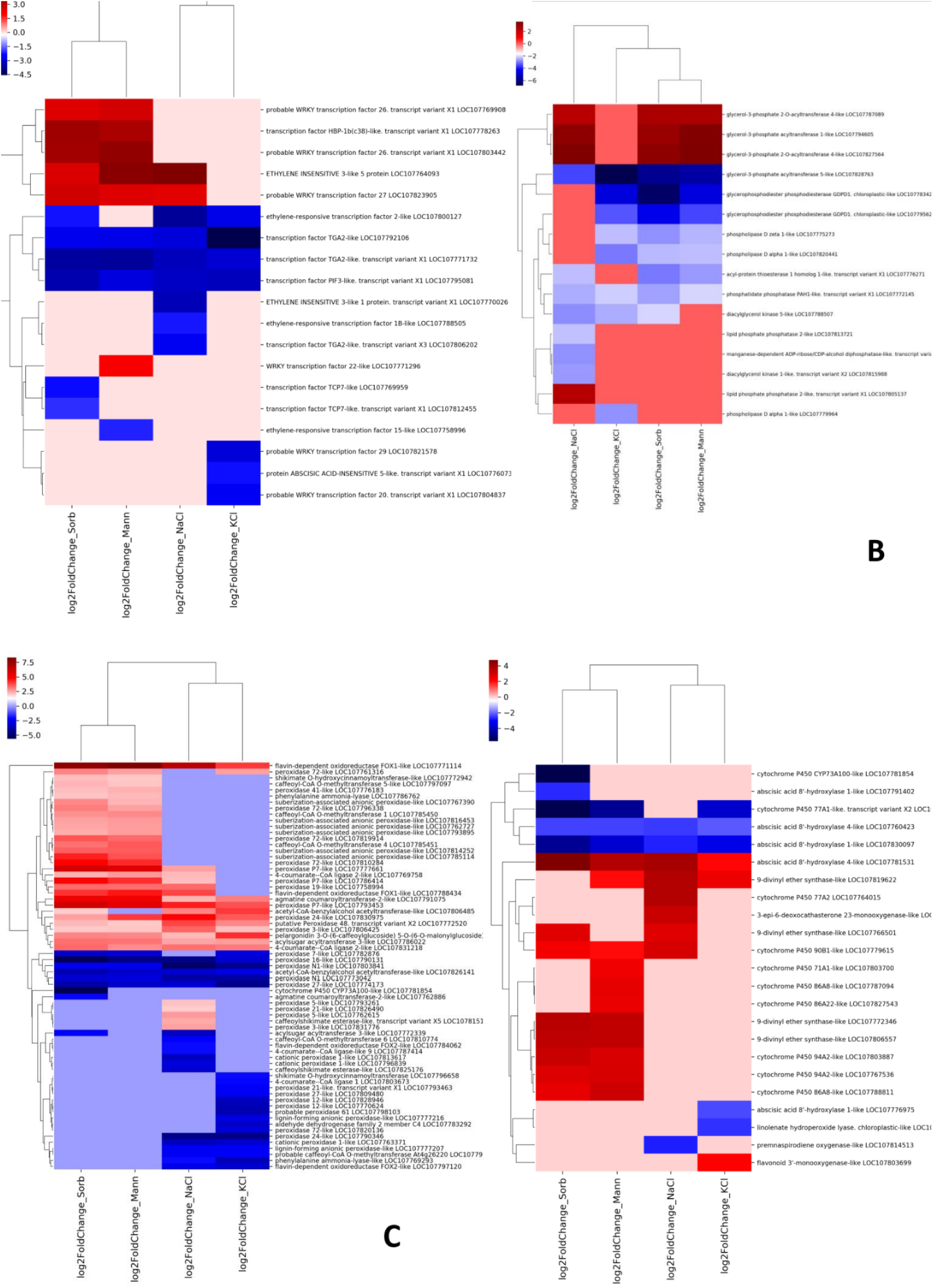
Heatmaps of log2 fold changes in gene expression. Genes derived from Nicotiana tabacum KEGG pathways: A) 03000 Transcription factors; B) 00564 Glycerophospholipid metabolism; C) 00940 Phenylpropanoid biosynthesis; D) 00199 Cytochrome P450

**Figure 6.**
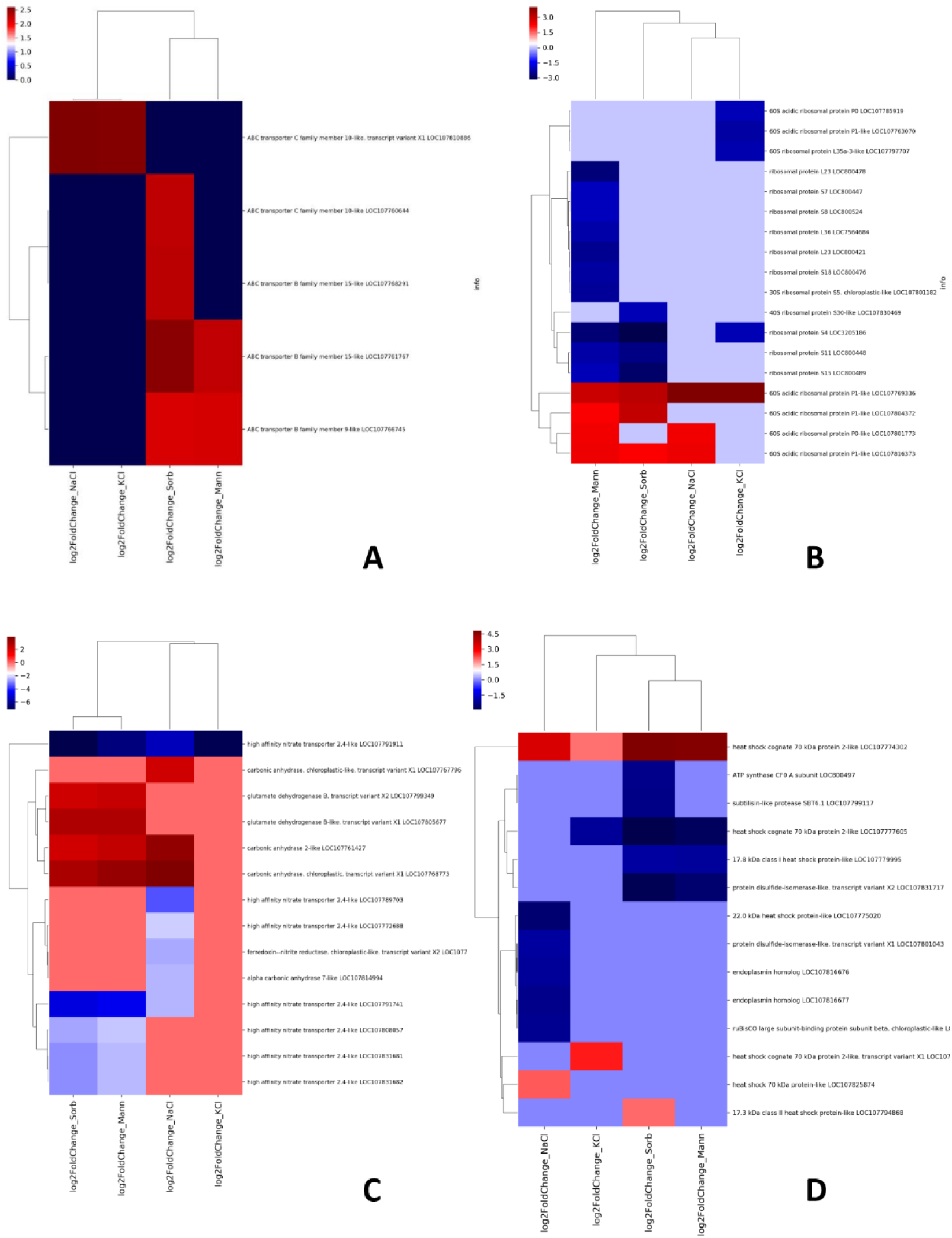
Heatmaps of log2 fold changes in gene expression. Genes derived from Nicotiana tabacum KEGG pathways: A) 02010 ABC transporters; B) 03010 Ribosome; C) 00910 Nitrogen metabolism; D) 03110 Chaperones and folding catalysts

SOS1 is up to date, the only known plants plasma membrane antiporter that provide Na^+^/H^+^ exchange. SOS1 expression provide tolerance to high salinity, and expression was boosted in response to salt stress as well^62^. Interestingly, SOS1 homologs from *N. tabacum* were not significantly up- or downregulated (data not shown). Similarly, expression of homolog genes encoding K^+^ channels (HAK5 (107832541), KT-1, GORK (LOC107781551)) - was also found to be not significantly shifted (data not shown). Previously, changes in expression of genes involved in monovalent cations efflux was shown as importance in adaptation of extremophyte *Schrienkella parvula,* which is more resilient than *A. thaliana* to high potassium. *S. parvula* boosts SOS1 and GORK expression in response to high potassium^27,63^. Also, in other in halophytes SOS1 genes expression also seem to be crucial for salt tolerance and adaptation^64^. Beside ionic channels, in plants ATP Binding Cassette (ABC) transporters genes are vastly represented and functionally versatile, also showing significance for stress tolerance. ABC transporters regulate hormones inter-intracellular transport and ions channels as well, and are important for stress tolerance^65,66^. Here, we identified also *N. tabacum* ABC transporters genes, that are specifically upregulated in high salts lines, and polyol lines and also two BY-2:Sorb specific ABC transporters (Fig. 6A ).

Analysis of DEGs that belong to known crucial genes in osmotic/salt stress response and in stress adaptation enable us to identify interesting similarities in gene families that were down- and upregulated. However, particular PP2Cs, SnRKs, PYR/PYLs, TFs altered their expression levels differently in all analyzed BY-2 lines. Such results suggest general similarities, but also phenomenon of osmotica-type specific response in adapted BY-2 cells. Interestingly, MAPK related pathways, as well as translation and ribosome related genes were more strongly refined in BY-2:Mann/Sorb, than BY-2:NaCl/KCl.

### Differences in BY-2 cell walls genes

Plants cell walls are cells interfaces that often first contact with factor that induce stress to cells. In recent years role of cell wall integrity surveillance system was revealed in perception of stress and adaptation process^20^. Despite previous results on composition of adapted cell walls BY-2^32,33^, there was no data of genes regulating stress perception and remodeling of plasma membrane-cell wall continuum, like malectin-like receptor protein kinase FERONIA, as well as, RALFs peptides (RAPID ALKALINIZATION FACTORs), and leucine-rich extensins (LRXs)^67–69^. Expansins, extensins and xyloglucan endotransglucosylase genes shows altered expression in salt-tolerance quinoa, and correlate with cell walls reorganization^14^. Despite various osmotica and BY-2 morphologies, here, only few of expansins, extensins and xyloglucan endotransglucosylase genes altered their expression, when compared to control BY-2 (Fig. 7). Moreover, the biggest changes concerned same genes for all lines.

**Figure 7.**
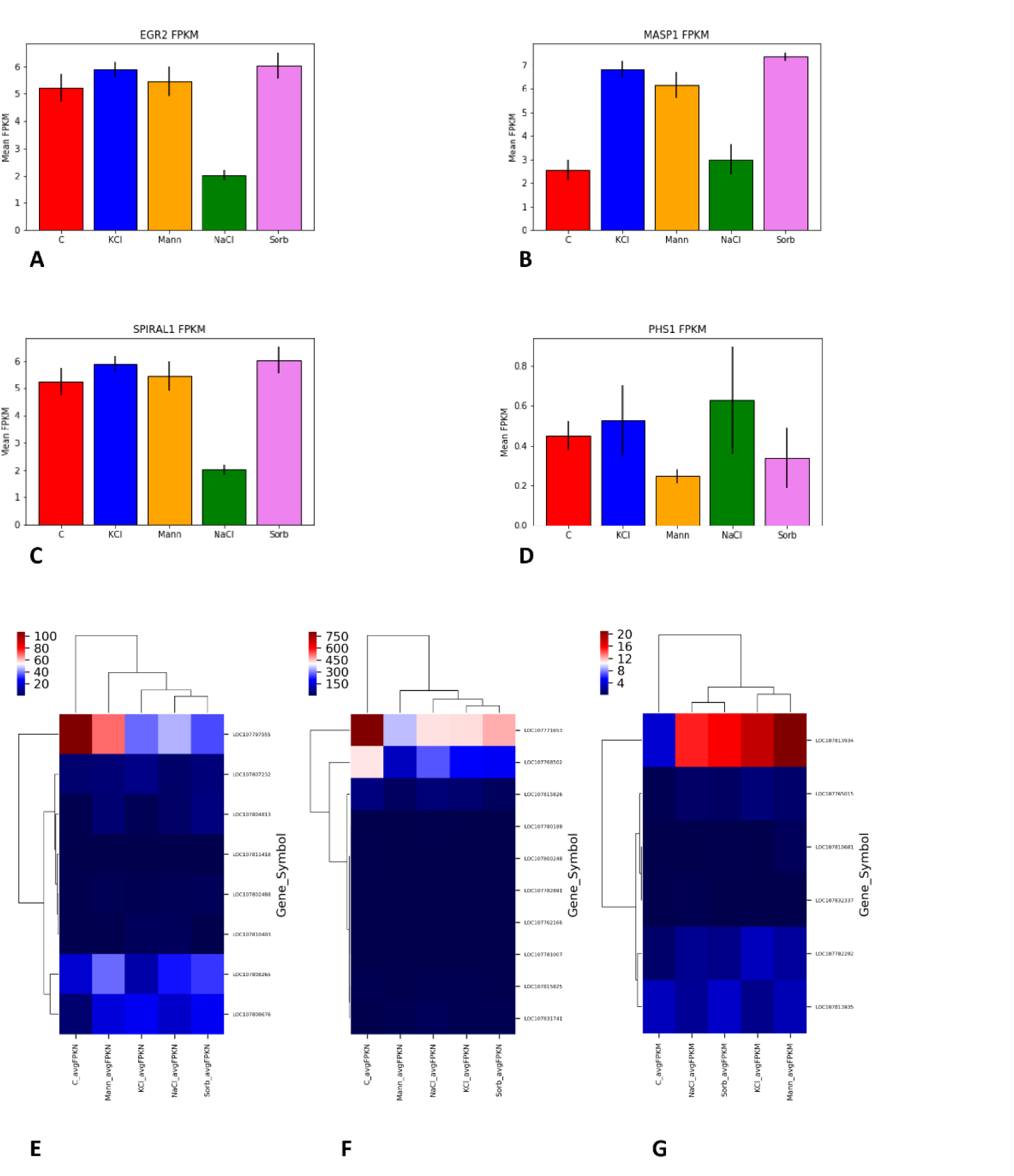
Expression level of A)EGR2; B) MASP1; C) SPIRAL1; D) PHS1 in N. tabacum BY-2 lines; E) expansins expression in FPKM; F) extensins expression in FPKM; G) xyloglucan endotransglucosylase in FPKM

Plants cell walls remodeling strongly dependent on cortical microtubules, cellulose synthase complex, and adjacent proteins. The microtubule (MT) interacting proteins were reported as involved in salt stress response, like PROPYZAMIDE HYPER-SENSITIVE1 (PHS1), that phosphorylates and inhibits microtubule polymerization, as MT stabilizing factor SPIRAL1^70,71^. PHS1 homolog (LOC107762514) exhibit only slight decreased in expression and only in BY-2:Mann and BY-2:Sorb. Oppositely, SPIRAL1 homolog (LOC107787772) was significantly downregulated, but only in BY-2:NaCl (Figure 5 CD). Previous results show that Arabidopsis root’s meristem subjected to drought stress is characterized by contradict expression and action of EGR2 and MASP1 on microtubule stability^72^. Here, consistently, BY-2:NaCl shows slightly distinct expression patterns of EGR2 (LOC107824506) and its downstream effector MASP1 (LOC107806103), than the other BY-2 lines (Figure AB). Such BY-2:NaCl specific expression pattern of EGR2/MASP1 might help BY-2 cells to stabilize microtubules and maintain cells division rate in environment with lowered water activity.

## Summary

Here we presented *Nicotiana tabacum* BY-2 suspension cell lines that provide useful model in investigation of plants cells subjected to long-term osmotic and salt stress. Utilization of few stress adapted lines would allow us to elucidate simultaneous impact of osmotic and ionic stress factors to particular cells adaptations.

High NaCl induce the most prominent transcriptional reprogramming in BY-2 suspension cells. Moreover, DEGs responsed to high potassium concentration have been identified and compared to DEGs from high NaCl and high mannitol/sorbitol cell lines. Interestingly, utilized various osmotica induce partially overlapping changes in expression of genes that belong to ABA responsive MAPK pathway, what was shown for *N. tabacum* treated with ABA and NaCl^73^.

Our results suggest importance of refined genetic reprograming required to survive and adapt to various osmotic stress in long term conditions. It was shown that in *A. thaliana* transcriptional reprogramming after salt, osmotic, and cold stress, there was less overlapping DEGs after 27 h, than after 3 h^2^. In more in-depth studies, by using combinations of cold, salinity, high light, heat, or flagellin, it was shown that ∼60% of transcriptional responses was non predicted when based on the response to single stresses alone^74^. Also, revealing the transcriptional response to salt, mannitol, heat, and combinations of them, showed that combined treatment induce uncomprehensive response with single stresses alone^75^. Recent data and analysis suggest severe decline of plats growth and survival due to combined stress conditions^76,77^. Here, transcriptomes are sequenced from long term, various osmotica adapted suspension cultures. Osmoticum repressed/induced genes partially overlap, with greater similarity within BY-2:Mann/Sorb and within BY-2:NaCl/KCl lines. Commonly in every line transcriptional reprogramming was massive and commonly drought related ABA, ethylene related genes expression was clearly shifted.

Gene expression data suggest that BY-2 lines have to permanently change their metabolism and signaling pathways, refine their stress and phytohormone metabolism in osmotica-type dependent manner.

In multiple metabolic pathways were found up- or downregulated genes. Hormones related signaling pathways and TFs play crucial roles in cells growth regulation, here we observe partially overlapping alterations of these genes expression. There is more overlapping DEGs within salts BY-2:NaCl/KCl and within polyols BY-2:Sorb/Mann, than between salt/polyol lines. The exact mechanism how metabolism and cells structure of BY-2 conform to endure such severe conditions, would require further research.

The presented adapted BY-2 suspension cell lines provide osmotica-specific interesting expression data in long-lasting transcriptional reprogramming. Moreover, such BY-2 model would allow further research on molecular mechanisms that enable plants cells to survive in ionic and osmotic stress conditions. Utilization of BY-2 lines enables research focus on similarities and discrepancies in gene expression, but also cells structure and physiology, when subjected to long-term various osmotic and high salt conditions.

